# Evolution of carbapenem resistance in *Acinetobacter baumannii* during a prolonged infection

**DOI:** 10.1101/228668

**Authors:** Jane Hawkey, David B. Ascher, Louise Judd, Ryan R. Wick, Xenia Kostoulias, Heather Cleland, Denis W. Spelman, Alex Padiglione, Anton Y. Peleg, Kathryn E. Holt

## Abstract

*Acinetobacter baumannii* is a common causative agent of hospital-acquired infections and a leading cause of infection in burns patients. Carbapenem resistant *A. baumannii* is considered a major public health threat and has been identified by the World Health Organization as the top priority organism requiring new antimicrobials. The most common mechanism for carbapenem resistance in *A. baumannii* is via horizontal acquisition of carbapenemase genes. In this study, we sampled 20 *A. baumannii* isolates from a patient with extensive burns, and characterized the evolution of carbapenem resistance over a 45-day period via Illumina and Oxford Nanopore sequencing. All isolates were multi-drug resistant, carrying two genomic islands that harboured several antibiotic resistance genes. Most isolates were genetically identical and represent a single founder genotype. We identified three novel non-synonymous substitutions associated with meropenem resistance: F136L and G288S in AdeB (part of the AdeABC efflux pump) associated with an increase in meropenem MIC to ≥8 μg/mL; and A515V in FtsI (PBP3, a penicillin-binding protein) associated with a further increase in MIC to 32 μg/mL. Structural modelling of AdeB and FtsI showed that these mutations affected their drug binding sites and revealed mechanisms for meropenem resistance. Notably, one of the *adeB* mutations arose prior to meropenem therapy but following ciprofloxacin therapy, suggesting exposure to one drug whose resistance is mediated by the efflux pump can induce collateral resistance to other drugs to which the bacteria has not yet been exposed.

**DATA SUMMARY:**

1. All raw genome sequences, including Illumina paired end short reads and Oxford Nanopore long reads, have been deposited in the SRA under project PRJNA396979. Individual accessions for each strain are provided in **Table S1**.
2. The annotated genome assembly for strain A2, the reference genome for the founder genotype, has been submitted to GenBank under accession CP024124 (chromosome) and CP024125 (plasmid).
3. Hybrid assemblies for strains A1, A3, A8, A13, A15, A17 and A20 are available in FigShare, doi: 10.4225/49/5987e14e9b530 (note they were not deposited in GenBank as they differ from A2 by only 1-4 SNPs as indicated).

**IMPACT STATEMENTS:** *Acinetobacter baumannii* is a highly drug resistant pathogen that is frequently found within intensive care units (ICUs) and especially impacts patients with severe burns. While several studies have examined the global population structure of *A. baumannii*, few have investigated within-host evolution of *A. baumannii* in direct response to antibiotic treatment in a single patient. Here, we analysed the genetic evolution of *A. baumannii* isolated from a patient with severe burns over the course of their stay in ICU. The *A. baumannii* population on this patient was highly drug resistant, carrying two distinct genomic islands encoding resistance to several antibiotics but not carbapenems. The bacterial population comprised four distinct subclades, two of which had evolved carbapenem resistance over the course of antibiotic treatment through novel mutations in genes associated with drug binding. One subclade was also transmitted to another patient on the ward. While carbapenem resistance is common in *A. baumannii*, this is generally attributed to horizontally transferred carbapenemase genes. These data provide evidence for carbapenem resistance arising *in vivo* via non-synonymous substitutions during a single infection episode, demonstrating carbapenem resistance can emerge in genetic isolation in response to exposure to carbapenems and other drugs.

## INTRODUCTION

*Acinetobacter baumannii* is a Gram negative bacteria that is amongst the six most common causes of multidrug resistant (MDR) hospital-acquired infections, a group collectively known as ESKAPE pathogens [1]. *A. baumannii* is a common causative agent of pneumonia, bacteremia, urinary tract infections, meningitis, and wound infections, especially in burns patients [2]. *A. baumannii* is intrinsically resistant to a broad range of antibiotics, however over the last three decades additional acquired resistance has emerged [3]. Acquired resistance to first line antibiotics, including aminoglycosides, sulfonamides and tetracycline arose in the 1970s, with most resistance genes residing on the chromosome within the resistance island AbaR, acquired independently by each *A. baumannii* lineage [4-9]. During the 1980s and 1990s, resistance to third-generation cephalosporins and fluoroquinolones emerged. Upregulation of the chromosomal beta-lactamase gene *ampC*, by ISAba1 or ISAba125, is the primary mechanism of cephalosporin resistance [10-12]. Fluoroquinolone resistance is primarily mediated through non-synonymous mutations in *gyrA* and *parC*, which interfere with drug binding [13-15]. Carbapenem resistance is becoming increasingly common, mainly through the acquisition of carbapenem-hydrolysing oxacillinase genes [3]. Carbapenem resistance has also been associated with changes in the expression of penicillin-binding proteins [16] or the AdeABC efflux pump [17,18], which can also impact resistance to other drugs such as fluoroquinolones [19]. The prevalence of carbapenem resistance in *A. baumannii* leaves only polymyxins (such as colistin) and tigecycline available for treatment. Resistance to these drugs has already been observed [3,20-22], and in 2017 the World Health Organization named carbapenem-resistant *Acinetobacter baumannii* as the top priority pathogen critically requiring research and development of new antibiotics [23].

The *A. baumannii* population primarily responsible for hospital infections in humans largely consists of two globally distributed clones, GC1 and GC2 [24]. Previous studies have begun to untangle the evolution of these clones [7,24], including the evolution of sub-clones specific to a particular hospital [25]. However, very few studies have investigated evolution of *A. baumannii* within individual patients. Within-host evolution has been studied in several other pathogens, including *Staphylococcus aureus*, *Helicobacter pylori* and *Burkholderia pseudomallei* [26-28]. One study retrospectively investigated within-host evolution of *A. baumannii* in patients from a single hospital system, revealing enrichment for mutations in genes responsible for antibiotic resistance or immune responses, with the *pmrAB* genes (linked to colistin resistance) being the most commonly mutated [29]. In addition, the regulatory genes *adeRS*, responsible for regulating the efflux pump AdeABC, were also found to be commonly mutated [29]. Another study compared the genomes of *A. baumannii* isolated from two patients pre-(sensitive) and post-(resistant) treatment with polymyxin B, and identified non-synonymous mutations in the *pmrB* gene in both cases (one GC1 and one GC2) [30]. These studies demonstrate mechanisms by which *A. baumannii* can adapt rapidly under positive selection from antimicrobial exposure during therapy with polymyxins. However, as both studies only examined pairs of isolates from each patient, they were unable to examine the diversification of the pathogen population during prolonged infection. Furthermore, while we have previously used genomics to unravel the mechanisms of evolution of carbapenem resistance at the level of an ICU [25], there are no published studies reporting within-host evolution of carbapenem resistance during an individual infection.

In this case study, we investigate the evolution of an *A. baumannii* strain within a single ICU patient, who had extensive burn wounds and developed a prolonged MDR *A. baumannii* infection. Treatment of this infection was complicated by the presence of resistance to most available drugs at the onset, followed by the emergence of resistance to meropenem after its therapeutic use.

Here, we compared the genomes of 20 isolates cultured from serial samples at nine time points spanning 68 days, including multiple samples taken at the same time from different body sites. We identified two distinct subpopulations of meropenem resistant strains harbouring non-synonymous substitutions in *adeB* and/or *ftsI*, and used structural modeling of the encoded proteins to demonstrate that these mutations can explain the observed loss of meropenem susceptibility.

## METHODS

### Bacterial isolates and antimicrobial susceptibility profiling

All *A. baumannii* strains cultured during the clinical care of a patient at The Alfred Hospital, Melbourne, Australia were included (n = 20). Single isolates from three other patients that were cared for in the same ward and during the same time as our index patient were also included. All bacterial isolates underwent susceptibility testing using automated methods (VITEK2), with confirmation of meropenem susceptibility and determination of MICs by E-tests as per manufacturers guidelines (Biomérieux, France).

### DNA extraction and sequencing

A. *baumannii* isolates were grown in LB overnight at 37°C with shaking. For the extraction of genomic DNA we first generated cell lysates using the Bacterial GenElute kit (Sigma) according to manufacturer’s instructions for Gram negative bacterial preparation. Cell lysates were then cleaned using the Qiagen DNeasy kit according to manufacturer’s instructions and DNA was eluted in nuclease free water. DNA sequencing libraries were prepared used the Nextera XT protocol, and 100 bp paired end sequencing was performed on each sample using the Illumina HiSeq 2500 at the Australian Genome Research Facility. Average insert size for each genome was 445 bp, with a mean read depth of 121x (**Table S1**). All reads were deposited in SRA under accessions listed in **Table S1**.

For long read sequencing, each isolate was grown overnight at 37°C on Luria Broth (LB) plates, and then single colonies were grown overnight at 37°C in LB. Bacterial cell pellets from 3.0 mL of broth culture were generated by centrifugation 15,000 *g* for 5 minutes. DNA was extracted from these pellets using Agencourt GenFind V2 (Beckman Coulter) with minor modifications, as described in [31]. No further purification or size selection is required, as this protocol generates high molecular weight gDNA that is free of small DNA contamination. Oxford Nanopore Technologies (ONT) sequencing libraries were prepared without shearing to maximise read length. Adapters were ligated to each library using the Nanopore 1D ligation sequencing kit (SQK-LSK108) with the native barcoding expansion kit (EXP-NBD103) and modifications as described in [31]. The resulting library contained 3,530 ng of DNA and was loaded onto an R9.4 flow cell. A MinION MK1b device (ONT) was used to perform the run using the NC_48Hr_Sequencing_Run_FLO-MIN106_SQK-LSK108 protocol, with 910 pore numbers available. At the completion of the run, all resulting fast5 files were transferred to a separate Linux server, where bases were called using ONT’s Albacore command line tool (v1.0.1), using barcode demultiplexing and fastq output. Adapter sequences were trimmed from the reads using Porechop (v0.2.0, https://github.com/rrwick/Porechop), with barcode demultiplexing, and only keeping reads where Albacore and Porechop agreed on the barcode bin, to prevent cross-barcode contamination. Mean read length was 9,148 bp and mean read depth per isolate was 59x (**Table S1**). The resulting demultiplexed, barcode trimmed reads were deposited in SRA under accessions listed in **Table S1**.

### Whole genome assembly

For the eight isolates for which both long (ONT) and short (Illumina) reads were generated, hybrid *de novo* assemblies were constructed using Unicycler v0.3.1 [32] with default settings. This resulted in complete assemblies with two circular replicons (one chromosome and one plasmid) for isolates A1, A2, A8, A13, A15, A17 and A20, and a single circular replicon (one chromosome) for isolate A3. For the remaining isolates, for which only Illumina-based short reads were available, *de novo* assemblies were generated using SPAdes v3.6.1 [33] with kmer sizes 21,33,45,57,69,81,93.All genome sequences (complete and draft) were annotated with RAST using default settings [34].

### Detection of resistance genes and multi-locus sequence type

Raw sequence reads were screened using SRST2 v0.2.0 [35] to identify (i) antibiotic resistance genes using the ARG-ANNOT database [36] and (ii) multi-locus sequence type (MLST) using the Oxford *A. baumannii* MLST scheme. Assemblies were also screened for antibiotic resistance genes using BLAST+ v2.6.0 [37], in order to determine the genetic context of resistance genes. Regions containing resistance genes were extracted from the assemblies and compared with known resistance islands in *A. baumannii* using BLAST+, and the comparisons visualized using ACT [38].

### Detection of variation between isolate genomes

#### Single nucleotide polymorphism (SNP) detection

All isolates were mapped to the completed A2 genome (i.e. the first strain isolated from the index patient) using RedDog v1beta10.3 (https://github.com/katholt/RedDog) with default parameters. Briefly, Illumina reads were mapped using Bowtie2 v2.2.9 [39] using the sensitive-local algorithm and a maximum insert length of 2000 (set with the x parameter). Variant sites (SNPs and indels) were called using SAMtools v1.3.1 [40]. Isolate A22 was identified as an outgroup, with only 87% coverage of the reference genome (compared to ≥99% for all other genomes). This isolate contained 103,519 SNPs, and was a different sequence type (ST) to the other strains, so was excluded from further analysis. Amongst the remaining 22 isolates, a total of 12 SNPs were identified.

#### Insertion sequence (IS) variation

ISSaga [41] was used to screen the A2 genome sequence for all known insertion sequences (IS). IS detected in the chromosome were ISAba22, IS26, ISVsa3, ISAba26, IS1326 and ISAba1. ISMapper [42] was used to screen all Illumina read sets for evidence of variation in copy number and location of the six IS in the other 21 genomes, in comparison to the A2 reference genome. The depth cut-off was set to ≥20x (rather than the default value, ≥6x) to avoid spurious calls in these deeply sequenced genomes (87–149x). Visual inspection of all ISAba1 hits in all genomes was also undertaken to confirm the exact location and orientation of each insertion site. The ISMapper results were compared to the eight finished genome assemblies belonging to the main clone, and were found to be in complete agreement concerning the position of all IS insertions.

#### Pairwise comparison of completed genomes

The eight completed, hybrid genome assemblies were subjected to pairwise base-by-base comparison using the *diffseq* function in the EMBOSS package [43]. This analysis confirmed all SNPs and indels detected by RedDog, the ISAba1 variation detected by ISMapper, and additionally identified: the loss of two tRNA genes within A2, the loss of a 52 kbp phage region in A8, an additional 1 bp intergenic insertion in A18 and A20, and a 1 bp insertion within Bm3R1 in A2, relative to all other genomes.

#### Structural modeling of mutations in AdeB and FtsI associated with meropenem resistance

Molecular models of AdeB and FtsI were generated using Modeller [44] and Macro Model (Schrodinger, New York, NY) using the X-ray crystal structures of homologous membrane pumps (PDB IDs: 1OY8, 2V50, and 4MT1; sequence identity 45-60%) and related penicillin binding proteins (PDB IDs: 3EQU, 3OC2, 3PBQ, 3PBN, 3UE3, 4BPJ, and 5ENS; sequence identity 43-87%), respectively. The models were then minimized using the MMF94s forcefield in Sybyl-X 2.1.1 (Certara L.P., St Louis, MO), with the final structures having more than 95% of residues in the allowed region of a Ramachandran plot. Meropenem was docked into the models using Glide (Schrodinger), and the position of the ligands in available crystal structures used to guide placement. The quality of the models was confirmed with Verify3D (data not shown). Model structures were examined using Arpeggio [45] and PyMOL.

The structural consequences of the missense variants were analysed to account for all the potential effects of the mutations [46].The effects of the mutations upon the stability of the proteins was predicted using SDM [47], mCSM-Stability [48] and DUET [49]. The effect of the mutations upon the binding affinity for meropenem were predicted using mCSM-Lig [50-52]. These computational approaches represent the wild-type residues structural and chemical environment of a residue as a graph-based signature in order to determine the change upon mutation in Gibb’s Free Energy of stability or binding.

## RESULTS AND DISCUSSION

During a 3-month period in 2013, 20 MDR *A. baumannii* isolates (A2–A20, A22) were cultured from swabs collected from a single patient (Patient 1) with extensive burns who was being treated in the ICU of a tertiary care hospital in Melbourne, Australia. The *A. baumannii* were isolated from swabs taken from several different sites across the body, sampled at nine time points (Figure 1b, Figure 2a). Three additional *A. baumannii* isolates (A1, A21 and A23), cultured from wound swabs from three other patients admitted to the same ICU ward within a month of the index patient's stay, were included in the study for comparison.

**Figure 1.**
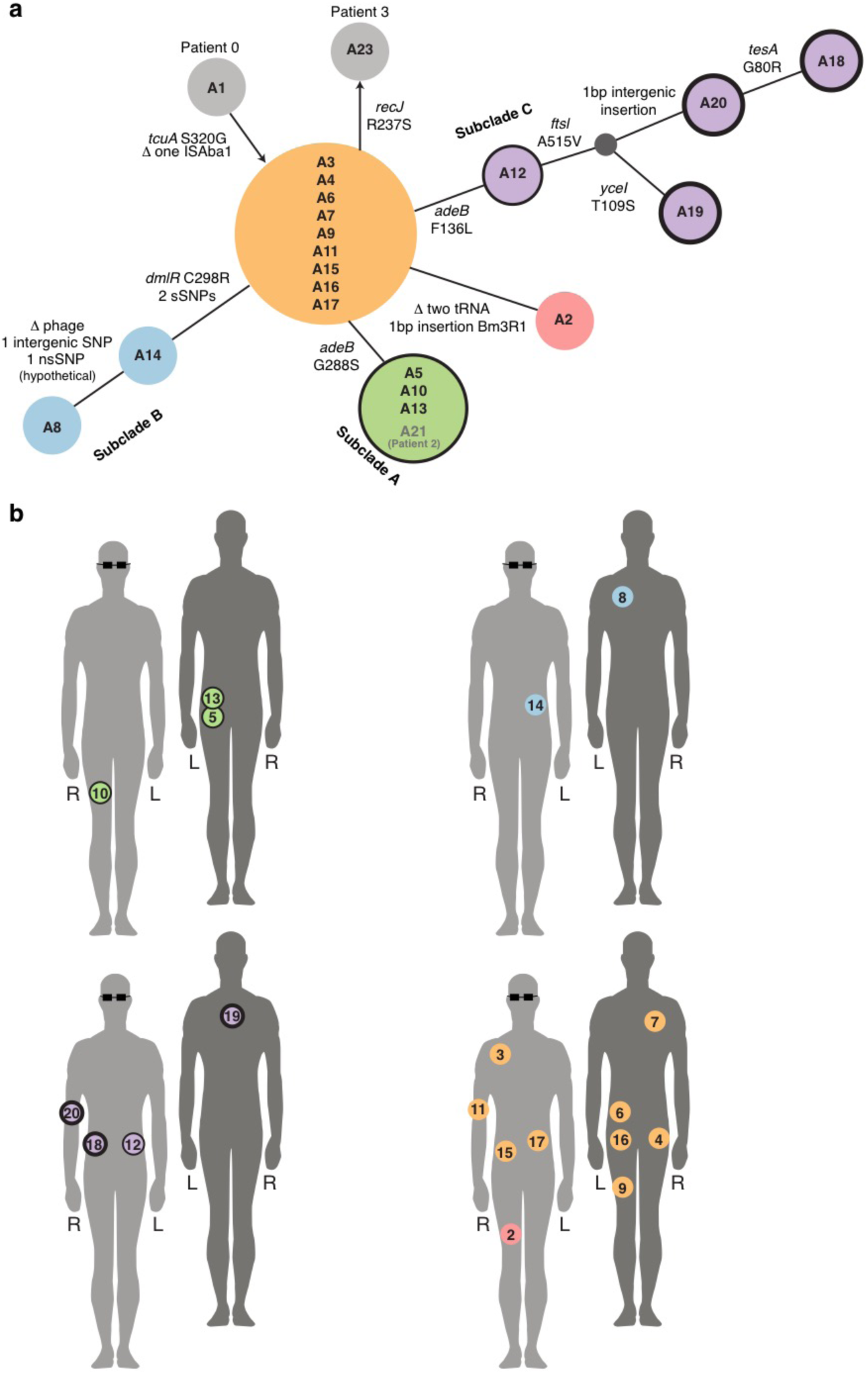
Population structure of *A. baumannii* and sample locations. **a**, Minimum spanning tree illustrating the population structure of all isolates, based on all genome variation detected. Circles indicate isolates whose genomes are genetically identical, with circle size relative to the number of isolates. Circles are coloured to indicate the subclades referred to in text; grey circles indicate isolates from other patients. Annotations above connecting lines indicate genome variation detected; non-synonymous SNPs in protein-coding genes are labeled with the effect on the amino acid sequence; synonymous SNPs (sSNPs) and intergenic SNPs are enumerated. Outlines around circles are indicative of meropenem MIC: no outline, MIC <8 μg/mL; thin outline, MIC ≥8 μg/mL; thick outline, MIC ≥ 32 μg/mL. **b**, Body map showing swab locations for samples of each lineage collected from Patient 1. Light grey with glasses, front; dark grey, back. Isolates are numbered in the order the swabs were collected. Circle outlines indicate meropenem resistance, as in **a**.

**Figure 2.**
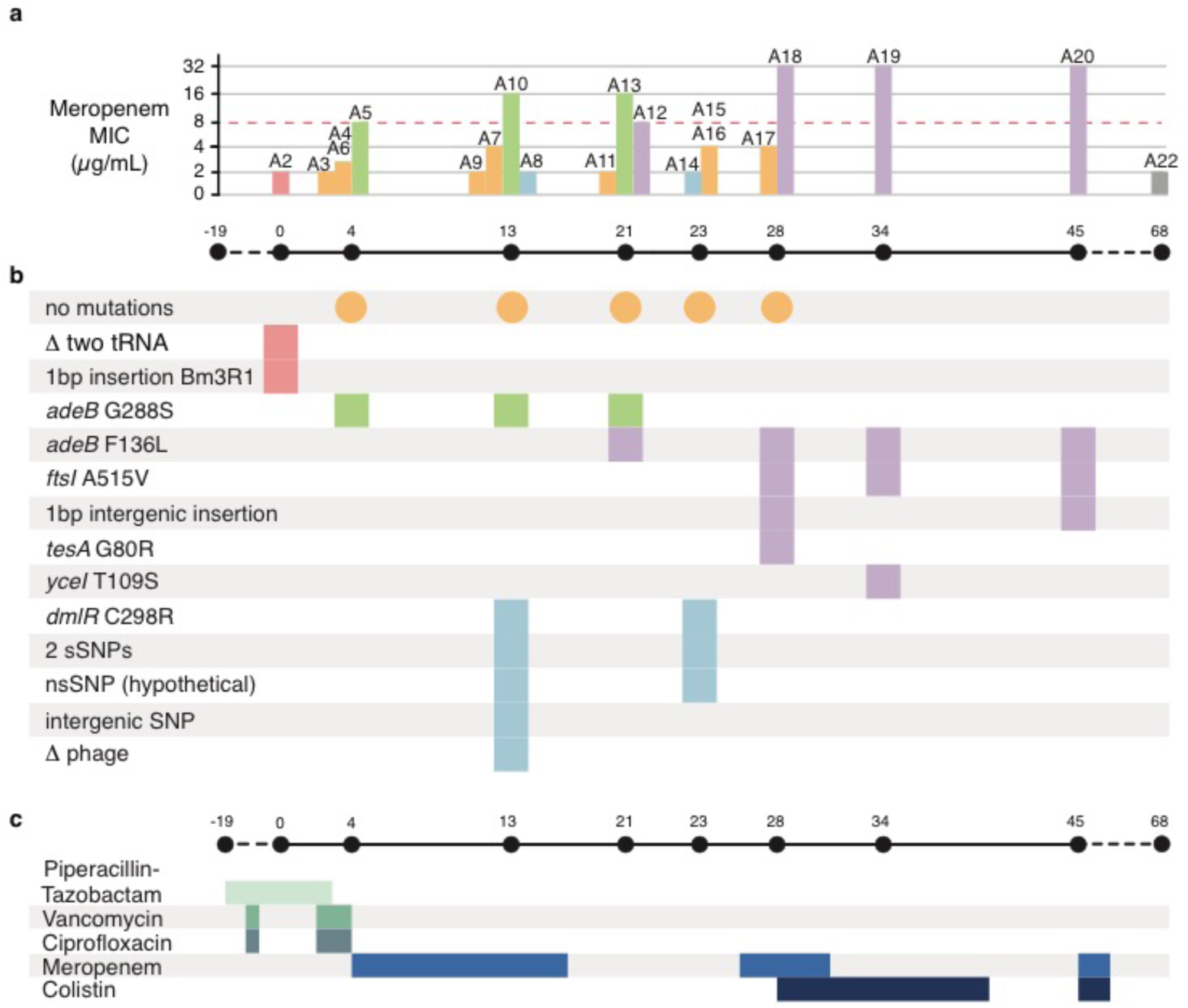
Infection timeline for Patient 1. **a**, Bar chart illustrating meropenem MIC for each isolate, with timeline below showing sample collection dates (relative to day 0 = first isolation from Patient 1). Bars are coloured by the genetic subclade, as defined in **Figure 1a**. Dashed red line shows the MIC=8 μg/mL, the EUCAST threshold for defining resistance. **b**, Temporal map of mutations identified in each isolate's genome, coloured by subclade. Orange circles indicate the dates on which the founder genotype was isolated; all genetic variation in other isolates are shown relative to this founder genotype. The delta symbol indicates deletion. **c**, Timeline of antibiotic treatment. Coloured bars show the length of time Patient 1 was treated with each antibiotic during their stay in the ICU.

All 23 *A. baumannii* isolates were sequenced using the Illumina platform. MLST and SNP analysis indicated that 22 of the isolates were GC2 (sequence type (ST) 2 according to the Institut Pasteur 7-locus MLST scheme) and formed a single clonal lineage with a maximum pairwise distance of eight SNPs (Figure 1a). A22 was a distinct strain of *A. baumannii*, separated from the main clonal lineage by >100,000 SNPs and belonged to a novel ST. This was the final isolate from Patient 1, cultured three weeks after the last GC2 isolate, and likely represents secondary infection with a novel strain.

### Variation amongst the 22 clonal isolates

The complete annotated genome sequence of strain A2, the first isolate cultured from Patient 1, was used as a reference for detailed comparative analyses. A total of twelve SNPs were detected amongst the 22 clonal GC2 isolates (9/12 non-synonymous, 2/12 synonymous, 1/12 intergenic; Figure 1a, Figure 2c). All isolates except A3 carried identical copies of the 110 kbp plasmid, confirmed by read mapping and assembly. Six different IS were present in the chromosome of A2, at a total of 37 unique sites: ISAba1 (23 sites), IS26 (8 sites), ISAba22 (3 sites), ISAba26 (1 site), ISVsa3 (1 site) and IS*1326* (1 site). All but one of the IS insertions present in A2 were conserved across all isolates. The exception was a single ISAba1 insertion between two genes encoding hypothetical proteins (coordinates 1,871,431–1,872,182 in A2), which was absent in A1 (the earliest isolate, from Patient 0), but present in all other genomes.

To facilitate high-resolution analysis of intra-host evolution, eight of the clonal GC2 isolates from Patient 1, and the index case from Patient 0, that displayed distinct genotypes and AMR phenotypes (strains A1, A2, A3, A5, A8, A15, A17 and A20) were selected for long read sequencing. The combined short and long read sequence data were used to assemble complete genomes for each of these isolates, which each consisted of a 3.9 Mbp chromosome and identical copies of a 110 kbp plasmid (with the exception of A3, which lacked any evidence of plasmid sequence in short or long reads). Comparison of the completed genomes to sequences in GenBank via BLAST searches revealed that the chromosome was most closely related to *A. baumannii* GC2 strain XH856 (isolated in 2010 in China; accession CP014541, 99% identity and 95% coverage), and the plasmid was similar to the previously described plasmid pABTJ2 (accession CP004359, 99% identity and 100% coverage) sequenced from *A. baumannii* GC2 strain MDR-TJ (isolated in China, year not reported) [53].

Base-by-base comparison between the eight finished genomes confirmed the SNP and IS variation data inferred from short reads, and additionally detected a single base pair intergenic insertion in A18 and A20, a single base pair insertion in Bm3R1 (a helix-turn-helix transcriptional repressor) in A2, the loss of two adjacent tRNA genes (tRNA-Trp and tRNA-Leu) from A2, and the loss of a 52 kbp phage region in A8 (**Figure 1a, Figure 2b**).

Nearly half (9/19) of the clonal isolates from Patient 1 formed a group of genetically indistinguishable strains (orange, **Figure 1a**) that were isolated from multiple body sites between days 4 - 28 (**Figure 1b**, **Figure 2a**). This dominant genotype occupied a centroid position in the minimum spanning tree of all strains from Patient 1, suggesting it likely represents the founder genotype in this patient. The isolates from the three other ICU patients, which were collected within one month before or after Patient 1’s infection, all belonged to the same clone and differed from the founder genotype by no more than one SNP (**Figure 1a**).

The remaining isolates from Patient 1 formed four related subclades each characterized by a unique set of differences from the founder genotype. Subclade A (n=3, green, **Figure 1**) was characterized by a single SNP and detected only on swabs taken from the left buttock and right anterior thigh area, early on in the series (day 4 - 21). Subclade B (n=2, blue**, Figure 1**) was detected on the left upper back and left flank in the middle of the series (day 13, day 23). Subclade C (n=4, purple, **Figure 1**) emerged via a stepwise series of mutations that accumulated later in the series. The initial SNP was first detected on the left trunk (day 21), and strains sharing this SNP plus additional mutations were subsequently detected on the right trunk (day 28), back (day 34) and right arm (day 45). The fourth subclade (D) was represented by a single isolate, A2, cultured from the right medial thigh on day 0.

To estimate the rate of within-host evolution occurring during the prolonged infection of Patient 1, we calculated for each isolate: (a) the temporal distance (in days) since the first detection of the clone in the patient (day 0), and (b) the genetic distance (in substitutions) from the founder genotype; and fit a linear regression to estimate the rate of substitutions per day (**Figure 3**). The fitted slope was 0.065, corresponding to a rate of ^~^1 SNP per fifteen days, or ^~^24 SNPs per year. This is higher than mutation rates previously reported, which were ^~^10 SNPs per year for GC2 *A. baumannii,* or ^~^5 SNPs per year in GC1 [7,25]. However, both studies examined *A. baumannii* populations that had been evolving for a significantly longer time period (10 years and 30 years, respectively), whereas the isolates considered here had only been evolving for ^~^45 days. It is known that slower substitution rates decay over time, as it takes time for purifying selection to act [54,55]. The relationship between substitution rate estimates for bacteria, and the sampling time (duration between first and last sample) of the isolate collection from which they were estimated, has recently been modeled using a decay function [56]. Using that decay function, the expected substitution rate for the 45-day period of evolution observed here is 7.3x10^-5^ substitutions site^-1^ year^-1^ (95% CI: 4x10^-5^ – 1.9x10^-4^), which is higher than the rate we measured here of 6.1x10^-6^ substitutions site^-1^ year^-1^ (95% CI: 2.9x10^-6^ – 9.2x10^-6^).

**Figure 3.**
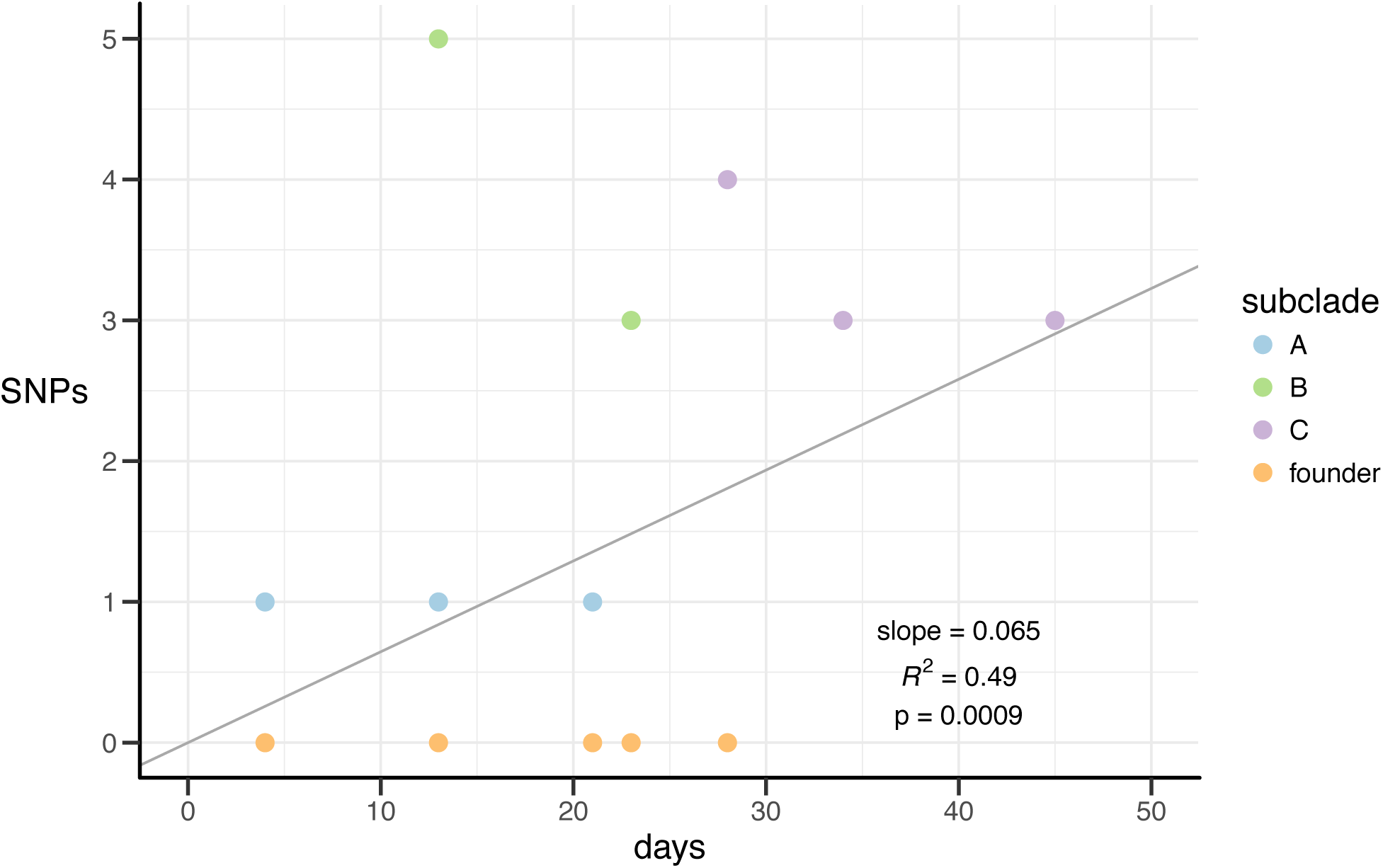
Estimated mutation rate for founder genotype and subclade A, B and C isolates from Patient 1. The number of SNPs observed in each isolate compared to the founder genotype (y-axis) is plotted against the number of days since the first isolate was collected from Patient 1.

### Evolution of antibiotic resistance

All *A. baumannii* isolates cultured from Patient 1 were MDR, displaying resistance to aminoglycosides, fluoroquinolones, trimethoprim-sulfamethoxazole, third- and fourth-generation cephalosporins (ceftazidime and cefepime) and carboxypenicillins. Variation was observed in sensitivity to meropenem (a carbapenem) (see **Figure 2** and detailed below).

All antibiotic resistance phenotypes besides meropenem could be explained by acquired resistance genes that were conserved in all the clonal (GC2) genomes. A variant of the AbGRI1-1 genomic island was present within the *comM* gene [57], encoding resistance to streptomycin (*strAB*), sulfonamides (*sul2*) and tetracyclines (*tetB*, *tetR(A)*). A variant of the resistance island AbGRI2-0 was also present [58]. This second resistance island included a partial mercury resistance operon (*mer*), and resistance genes for aminoglycosides (*aph3*’’*Ia*, *aadA1*, *aac3-I*), sulfonamides (*sul1*) and ampicillin (*bla*_TEM-1_). In addition to acquired resistance genes, all isolates also harboured an ISAba1 insertion site upstream of the intrinsic efflux pump *ampC*, which is known to generate resistance to third-generation cephalosporins [10]. The *bla*_*OXA*-66_ gene was also present within the chromosome. While the encoded Oxa-66 protein can have intrinsic carbapenemase activity, upregulation via IS elements or structural changes via non-synonymous mutations within the protein are required for full carbapenem resistance, neither of which were identified in these genomes (and all had identical *bla*_*OXA*_-_66_ gene sequences and promoter regions).

The majority of isolates, including all with the founder genotype (orange, **Figures 1-2**), displayed meropenem MIC ≤ 4 μg/mL (**Table S1**). Reduced sensitivity to meropenem was first observed in subclade A, which differed from the founder genotype by a non-synonymous substitution (G288S) in AdeB (green, **Figures 1-2**). These isolates displayed an increased MIC for meropenem (≥8 μg/mL). Subclade C isolates also displayed increased meropenem MICs and carried a different substitution (F136L) in AdeB (purple, **Figures 1-2**). Isolate A12 differed from the founder genotype by this SNP alone, and displayed an MIC of 8 μg/mL. The later isolates A18, A19, A20 (days 28, 34 and 45, see **Figures 1-2**) carried the AdeB F136L mutation and an additional substitution in the penicillin-binding protein (PBP3) FtsI (A515V), and displayed high-level resistance to meropenem (MIC ≥32 μg/mL).

The schedule of antimicrobial therapy supplied to Patient 1 is shown in **Figure 2c** and included two courses of meropenem. The meropenem resistant subclade A emerged prior to the initiation of meropenem treatment and persisted after its completion; this is potentially because the AdeB efflux pump can also transfer ciprofloxacin [19,59] to which the patient had been exposed prior to first detection of any AdeB mutations (**Figure 2**). The first meropenem resistant subclade C isolate, A12, was also collected after the completion of the first course of meropenem. However, two meropenem sensitive isolates were subsequently recovered, and a second course of meropenem treatment was initiated. A few days later, subclade C isolates with the additional *ftsI* mutation and high-level meropenem resistance (MIC >32 μg/mL) were recovered, prompting the introduction of colistin for treatment, with subsequent resolution of infection.

### Structural basis for meropenem resistance

It is striking that both subclades displaying increased MICs for meropenem carried non-synonymous mutations in the same protein, AdeB. The AdeABC efflux pump system has previously been implicated in carbapenem resistance, through the upregulation of the trans-membrane protein AdeB [3,60]. Here, we observed resistance (MIC ≥8 μg/mL) in all strains carrying either a G288S (subclade A, green) or an F136L (subclade C, purple) mutation in AdeB. Structural modelling of AdeB (see **Methods**) showed that the residues affected by these two mutations are located within the intracellular domain of the protein, which is responsible for the recognition of compounds to be passed to the outer membrane protein AdeC. Both mutations were within 5 Å of the bound meropenem in the model (**Figure 4a**). Both mutations were predicted to mildly destabilise the local structure (ΔΔG_Stability_^G288S^ = −0.62 ± 0.15 Kcal/mol and ΔΔG_Stability_^F136L^ = −0.91 ± 0.14 Kcal/mol), opening the drug binding site and pore, allowing meropenem to be more effectively exported.

**Figure 4.**
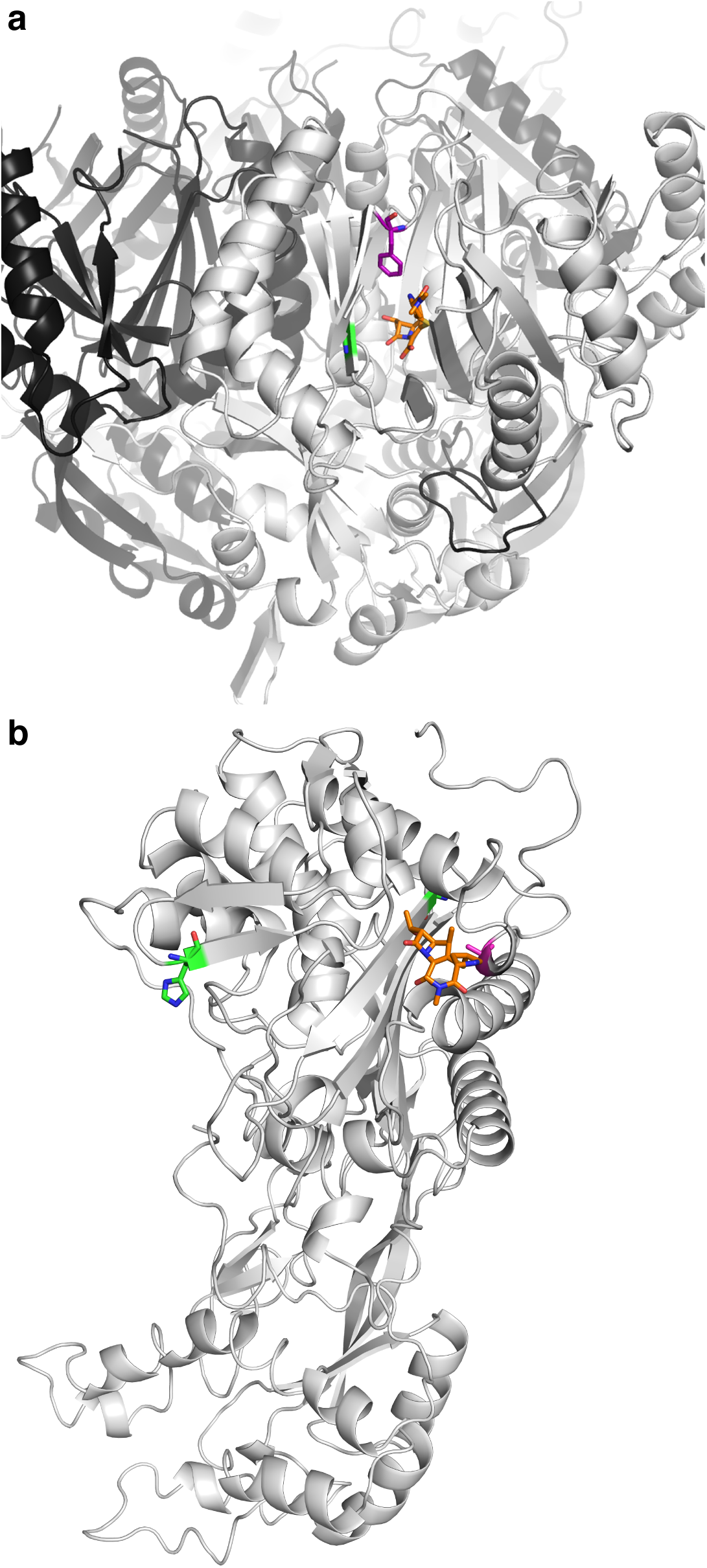
Meropenem bound to AdeB and FtsI. a, AdeB. The substitutions G288S (subclade A, green sticks) and F136L (subclade C, purple sticks) are predicted to facilitate binding and export of meropenem, shown here as the orange molecule. **b, FtsI**. The substitution A515V (magenta sticks) is located in close proximity to the binding site for meropenem (orange molecule) and is predicted to increase the binding affinity and sequestration of the drug. Other variants G523V and H370Y (green sticks), previously observed in carbapenem sensitive strains, are predicted to either reduce or have little effect on the binding affinity.

High level resistance (MIC ≥32 μg/mL) was observed only in the three subclade C strains that carried the FtsI A515V mutation in addition to AdeB-F136L. FtsI is a class B penicillin-binding protein, associated with cell division [61]. Two mutations (G523V, H370Y) have been reported previously in this protein, but they showed no evidence of contributing to carbapenem resistance [61]. Here we modelled these mutations and the A515V mutation present in our meropenem resistant isolates. The G523V mutation is located approximately 9 Å away from the meropenem binding site, and was predicted to greatly destabilise drug binding (-2.6 ± 0.05 log_10_(fold change in affinity)). By contrast, H370Y is located quite distal from the binding site (20 Å) and is predicted to have minimal effect on the affinity for meropenem (-0.03 ± 0.08 log_10_(fold change in affinity)). In our data, the mutation A515V is located close (7 Å) to the drug binding site of the protein (**Figure 4b**). Structural modelling indicated that mutation to valine would be well tolerated (mildly stabilising, ΔΔG_Stability_ = 0.25 Kcal/mol) and lead to a significant increase in the binding affinity for meropenem (0.16 ± 0.04 log_10_(fold change in affinity)), explaining the reduced effectiveness of meropenem against these strains.

## CONCLUSION

Here, we showed that the prolonged infection of Patient 1 was primarily due to spread of a founding *A. baumannii* genotype, acquired from Patient 0. Over the course of antibiotic treatment in Patient 1, the *A. baumannii* population diversified into three distinct subclades that were associated with time periods and specific spatial zones on the body of Patient 1, in addition to onward transmission to two other patients on the ward. During the evolution of this infection, two meropenem resistant lineages (MIC ≥8 μg/mL) emerged via non-synonymous substitutions in AdeB, with further increase in MIC (≥32 μg/mL) due to a non-synonymous substitution in FtsI. The majority of resistance to meropenem and other carbapenems in *A. baumannii* is due to acquisition of carbapenemases via horizontal gene transfer; indeed prior to this study, the upregulation of the AdeABC efflux pump was the only other known mechanism of meropenem resistance in *A. baumannii* [3,60,62,63].

Previous studies investigating *A. baumannii* evolution over short time periods have mostly focused on several patients within the same hospital system, with only a few samples taken per patient [29,64]. These studies indicated that reinfection with novel strains often occurred after treatment [64]. In our patient, only a single isolate taken at the very end of the series was a completely novel strain, diverged from the founder genotype by over 100,000 SNPs. The presence of this novel strain occurred three weeks after the core group of strains and following apparent resolution of the infection, and may have arisen due to treatment with colistin eradicating the founding strain population. However, at later time points, only single swabs were taken, and so this potentially underestimates the diversity of the population beyond the final course of antibiotic treatment.

These data highlight the therapeutic risks involved with treating high burden *A. baumannii* infections, especially in the context of prolonged and severe burn wounds. The phenotype-genotype relationships also highlight the importance of antibiotic selection pressure being able to drive resistance not only to the agent being used, but also to other antibiotics through shared resistance mechanisms. For example, the study suggests the activity of multidrug efflux pumps such as AdeABC may be increased through exposure to quinolones, with subsequent effects on meropenem susceptibility. The patient history also highlights the importance of cumulative mutations that can ensue after recurrent antibiotic exposures, culminating in extreme drug resistance, which supports the practice of taking into account prior antibiotic exposures when deciding on new empiric treatment of recurrent burn wound infections. The case exemplifies how *A. baumannii* is notorious for its rapid and efficient ability to genetically adapt to antibiotic selection pressures, both upregulating and downregulating resistance dependent on active antibiotic treatment regimens.

## ACKNOWLEDGEMENTS

KEH, AYP and DBA were supported by Fellowships from the NHMRC of Australia (APP1061409, APP1117940, APP1072476). DBA was also supported by the Jack Brockhoff Foundation (JBF 4186, 2016).

---

**Table S1: Isolates sequenced in this study.** Total number of reads, average chromosomal depth (Illumina and ONT), total number of sequenced bases (ONT only), mean insert size (Illumina only), NCBI accessions, meropenem MIC, patient number, day of collection, ST and clade for each isolate.

